# Spatiotemporal regulation by downstream genes of *Prok2* in olfactory system: from development to function

**DOI:** 10.1101/2024.10.31.621428

**Authors:** Bo-Ra Kim, Min-Seok Rha, Hyung-Ju Cho, Joo-Heon Yoon, Chang-Hoon Kim

## Abstract

Olfaction is important for the quality of life; however, in Kallmann syndrome (KS), defective development results in olfactory dysfunction. The mechanism underlying olfactory development, especially in the olfactory epithelium (OE), which detects olfactory signals, remains unclear. Mutations in *PROK2*, which encodes prokineticin-2, occur in approximately 9% of the KS patients with olfactory defects. In this study, we examined olfactory function and analyzed the causes of olfactory dysfunction based on spatiotemporal development and gene expression changes in *Prok2* knockout (KO) model mice with KS. Adult *Prok2* KO mice exhibited reduced ability to detect olfactory signals. Maturation of olfactory sensory neurons (OSNs) in the OE and formation of glomeruli in the olfactory bulb (OB) in adult *Prok2* KO mice were disrupted, thus causing olfactory dysfunction. Molecular analysis of *Prok2* KO mice during embryonic development revealed abnormal development of OB layers and reduced differentiation to mature OSNs in the OE, which caused defects in the entire olfactory system. Downstream signaling genes of *Prok2*, including intermediate filament genes and genes expressed in the putative OB, mediated olfactory system organization. Our findings reveal the role of *Prok2* in olfactory system organization, providing insights into the mechanisms through which olfactory development defects translate to olfactory function.

## Introduction

Olfaction is an essential sense for maintaining a good quality of life, daily life, and social interaction. Reduced olfactory function is an indicator of several diseases, including neurodegenerative disease (Doty 2017; Marin et al. 2018).

Olfactory dysfunction can result from various factors, including brain disorders such as brain tumors or traumatic brain injury (Proskynitopoulos et al. 2016; Li et al. 2023), inflammation-induced structural changes to the nasal cavity (Gudis and Soler 2016), neuronal degeneration caused by pathogen infection or neurogenerative disease (Murphy 2019; Tsukahara et al. 2023), and abnormal development due to genetic variation (Dodé and Rondard 2013; Topan et al. 2021). Kallmann syndrome (KS) is a developmental disease characterized by hypogonadotropic hypogonadism and congenital olfactory dysfunction (Rugarli 1999; Dodé and Rondard 2013; Forni and Wray 2015). It is primarily caused by the failure of hormone neurons to migrate into the brain during early development leading to subsequent collapse of reproductive axis formation and olfactory bulb (OB) construction (Schwanzel-Fukuda et al. 1994; Watanabe et al. 2009; Barraud et al. 2013; Taroc et al. 2017; Taroc et al. 2020a; Taroc et al. 2020b). Examples of hormone neurons include gonadotropin hormone-releasing hormone (GnRH) neurons or luteinizing hormone-releasing hormone neurons. Further, mutations of some genes such as *Anosomin-1* (*Kal-1*), *Prok2*, *Prokr2*, *Fgf8*, *Fgfr1*, *Gli3*, *Isl1*, *Sox10*, *Fezf1*, and *Sema3a* are associated with the migration of hormone neurons, which suggests that most of these genes regulate the formation of the nasal forebrain junction (NFJ) (Watanabe et al. 2009; Dodé and Rondard 2013; Taroc et al. 2020a; Taroc et al. 2020b).

Prokineticin 2 (Prok2) is a secretory protein that interacts with prokineticin receptor 2 (Prokr2) (Dodé et al. 2006; Sarfati et al. 2010; Dodé and Rondard 2013). *Prok2* gene comprises four exons (exons 1–4), in which exon 3 undergoes alternative splice processing (Dodé et al. 2006; Sarfati et al. 2010; Dodé and Rondard 2013). Among the different mutations, primarily missense mutations exons 1 and 2 have been reported in KS patient with olfactory dysfunction and/or hypogonadotropic hypogonadism. Notably, exon 1 includes the AVITGA motif, which is essential for the biological activity of the peptide and exon 2 includes the cysteine-rich region, which plays a role in protein folding (Dodé et al. 2006; Sarfati et al. 2010; Dodé and Rondard 2013). The overall frequency of *PROK2* or *PROKR2* mutations in KS patients are estimated to be approximately 9% (Sarfati et al. 2010; Dodé and Rondard 2013). *Prok2* knockout (KO) mice, which exhibit a deletion of the exon 2 allele, show hypoplasia of OB and reproductive system (Pitteloud et al. 2007). Additionally, Prok2/Prokr2 signaling is involved in the migration of GnRH from the vomeronasal organ (VNO) to the OB and in the migration of neuronal stem cells from the subventricular zone to the OB by interacting with Prokr2 (Ng et al. 2005; Pitteloud et al. 2007; Wen et al. 2019). However, the defects in the olfactory epithelium (OE), which detects olfactory stimuli, and the modulator of olfactory system formation during olfactory development remain unexplored. Here, we aimed to examine the changes in olfactory structure occurring during embryonic development as well as the genes regulated by *Prok2* to reveal the causal mechanism of olfactory dysfunction using *Prok2* KO mice as a KS model.

## Result

### *Prok2* KO mice with OB hypoplasia and olfactory dysfunction are a suitable model mimicking KS patients presenting with olfactory disorders

Consistent with OB hypoplasia in patients with KS (Sarfati et al. 2010; Wakabayashi et al. 2021; Bortolotto Felippe Trentin et al. 2023), *Prok2* KO mice exhibited a smaller OB than that of wild-type (WT) mice in the view of the medial-dissected head (dashed line in Fig. 1A). Although olfactory dysfunction has been reported in patients with KS (Wakabayashi et al. 2021; Bortolotto Felippe Trentin et al. 2023), the olfactory function of *Prok2* KO mice remains unclear. To evaluate the ability of *Prok2* KO mice to detect and recognize olfactory signals, these mice were subjected to an olfactory behavior test. The avoidance time of KO mice against trimethylthiazoline (TMT) was lower than that against water (Fig. 1B). Further, the avoidance time of KO mice against TMT was significantly lower than that of WT mice (KO mice mean ± SEM = -28.83 ± 10.85 s vs. WT mice mean ± SEM = 40.71 ± 11.82 s; p = 0.0010). These results indicated that *Prok2* KO mice have defective olfactory cognition.

**Figure 1.**
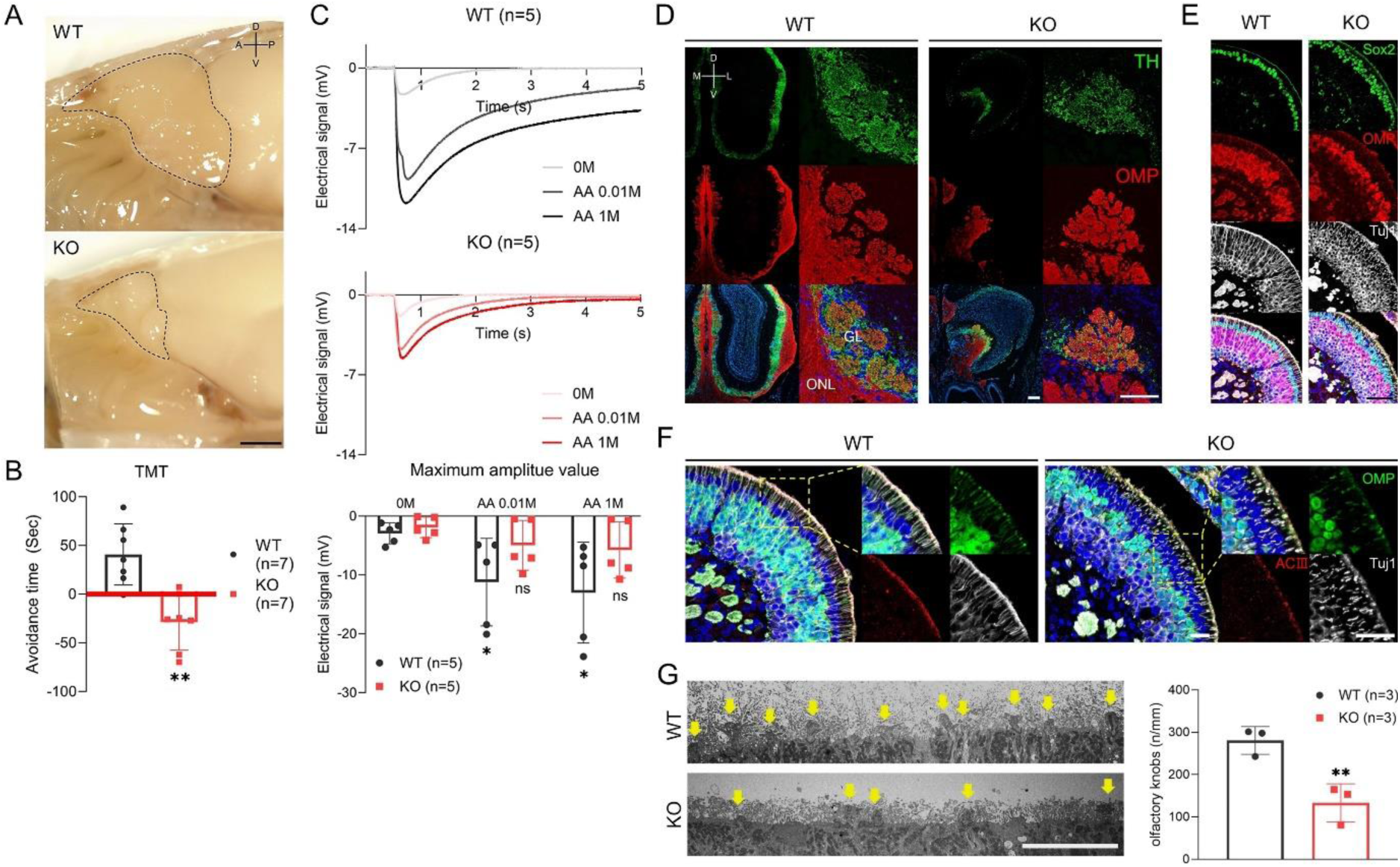
Olfactory bulb hypoplasia, olfactory dysfunction, and dysgenesis of the olfactory bulb, olfactory epithelium, and olfactory cilia in adult *Prok2* KO. (*A*) Olfactory bulb (OB) structure of WT and *Prok2* KO mice in the sagittal view. (*B*) Graph of the avoidance test. For avoidance time, p = 0.0010 for comparing the WT (*n* = 7) and *Prok2* KO mice (*n* = 7). Data are presented as mean ± SD. (*C*) Graphs of the time-dependent electrical signal for WT (*n* = 5) and *Prok2* KO mice (*n* = 5) and the graph of dose-dependent electrical signal of WT (*n* = 5) and *Prok2* KO mice (*n* = 5). In WT mice, the difference between electrical signal at 0 M and at 0.01 M has p = 0.0417, and that between electrical signal at 0 M and at 1 M has p = 0.0327. In *Prok2* KO mice, the difference between the electrical signal at 0 M and at 0.01 M has p = 0.3505, and that between the electrical signal at 0 M and at 1 M is p = 0.1301. Data presented as mean ± SD. (*D*-*H*) Immunostaining images, transmission electron microscopy (TEM) images, and a graph showing the number of olfactory knobs in WT and *Prok2* KO mice at 16 weeks. (*D*) Immunostaining of the olfactory bulb with α-TH and α-OMP antibodies. (*E*) Immunostaining of the olfactory epithelium with α-Sox2, α-OMP, and α-Tuj1 antibodies. (*F*) Immunostaining of the olfactory epithelium and olfactory cilia with α-OMP, α-ACⅢ, and α-Tuj1 antibodies. (*G*) Transmission electron microscopy (TEM) image of the olfactory cilia and a graph of the number of olfactory vesicles. P = 0.0015 between the number of olfactory knobs in WT (*n* = 3) and *Prok2* KO mice (*n* = 3). Data presented as mean ± SD. GL, glomerular layer; ONL, olfactory nerve layer; Sus, sustentacular cells; mOSN, mature olfactory sensory neuron. D, dorsal; V, ventral; A, anterior; P, posterior; M, medial; L, lateral. Scale bar, 1 cm in (*A*); 200 μm in (*D*) and (*E*); 100 μm in (*F*); and 10 μm in (*F*). For all panels, *n* indicates biologically independent repeats. Student’s *t*-test, p > 0.05 (ns), p < 0.05 (*), and p < 0.01 (**).

Olfactory cognition refers to the unified response of both central and peripheral olfactory systems. To identify whether the disruption of olfactory cognition in *Prok2* KO mice originated from the peripheral level in the olfactory system, the electrical signal against odor stimuli was measured in KO mice using an electro-olfactogram (EOG). The electrical signal against the stimuli of 0.01 M or 1 M amyl acetate in WT mice was significantly higher than that against the stimulus of a vehicle (0.01 M amyl acetate mean ± SEM = -11.21 ± 3.33 mV vs. vehicle mean ± SEM = -2.93 ± 0.79 mV, p = 0.0417; 1 M amyl acetate mean ± SEM = -13.00 ± 32.83 mV vs. vehicle, p = 0.0327) (Fig. 1C). In contrast, the electrical signal against the stimuli of 0.01 M or 1 M amyl acetate in KO mice was not significantly higher than that against the stimulus of a vehicle (0.01 M amyl acetate mean ± SEM = -4.96 ± 1.89 mV vs. vehicle mean ± SEM = -1.92 ± 0.78 mV, p = 0.3505; 1 M amyl acetate mean ± SEM = -5.76 ± 2.14 mV vs. vehicle, p = 0.1301). These results indicate that the dose-dependent response against odor stimuli is degraded in *Prok2* KO mice. Overall, the reduced ability of the peripheral olfactory system affects the disruption of olfactory cognition in *Prok2* KO mice. Thus, *Prok2* KO mice exhibit olfactory dysfunction similar to that in patients with KS.

### Olfactory structure dysgenesis in adult *Prok2* KO mice

Abnormalities in the OB and the olfactory function of Prok2 KO mice implicate dysgenesis in the olfactory system, which includes both OB and OE. Observation of the glomerular layer (GL) of OB showed that the radial structure of glomeruli was normally formed in WT mice (Fig. 1D). In contrast, only a few glomeruli were formed with interaction of TH^+^ periglomerular neurons and OMP^+^ olfactory axons in KO mice (Fig. 1D). In the OE of KO mice, the Sox2^+^ sustentacular layer was multi-layered. The OMP^+^ mature OSN layer was thinner and the OMP^−^Tuj1^+^ immature OSN layer was thicker than those in WT mice (Fig. 1E). Thus, the overall cellular composition of KO mice was disrupted. Higher magnification revealed that the ACⅢ^+^ or Tuj1^+^ olfactory cilia layer in KO mice was thinner than that in WT mice (Fig. 1F). Further, the number of olfactory knobs observed at a higher magnification using transmission electron microscopy (TEM) in KO mice (43.74 ± 23.67 n/mm) was significantly less than that in WT mice (280.57 ± 18.97 n/mm) (yellow arrows in Fig. 1G). Likewise, this relatively reduced number of olfactory knobs in KO mice was even observed in the OMP^+^ olfactory cilia layer (Fig. 1E). Thus, *Prok2* mediated GL formation in the OB, the composition of sustentacular cells and OSNs in the OE, differentiation of immature OSNs into mature OSNs, and the development of the olfactory cilia layer and olfactory knob.

### Expression pattern of *Prok2* and its receptor genes in the olfactory system from embryonic development to the adult stage

*Prok2* and *Prokr2* are expressed in the migratory route of hormone neurons from the VNO to the brain in the early embryonic stages, in the olfactory ventricle of the putative OB during the embryonic stages, and in most interneurons after birth (Ng et al. 2005; Pitteloud et al. 2007; Zhang et al. 2007; Wen et al. 2019). However, whether *Prokr1*, another receptor gene of *Prok2*, is expressed in the olfactory system and whether *Prok2* and its receptor genes are expressed in the OE remains unclear. Therefore, the expressions of *Prok2*, *Prokr2*, and *Prokr1* were assessed at E11.5, E14.5, E18.5, and 16 weeks. *Prokr1* was expressed in the OE and OB during embryonic development and only in the OB at 16 weeks (Fig. 2). This expression pattern was similar to that of *Prok2.* However, *Prokr1* was not expressed in the migratory route of hormone neurons, whereas Prok2 and Prokr2 were expressed (arrows in Fig. 2). Hence, Prok2/Prokr1 signaling is involved in the olfactory system, similar to Prok2/Prokr2 signaling. Further, *Prok2*, *Prokr2*, and *Pokr1* were all expressed in the OE during embryonic development. In contrast, they were almost absent in the OE at 16 weeks (Fig. 2). Thus, both Prok2/Prokr2 signaling and Prok2/Prokr1 signaling interact in OE, but only the former mediates the migration of hormone neurons.

**Figure 2.**
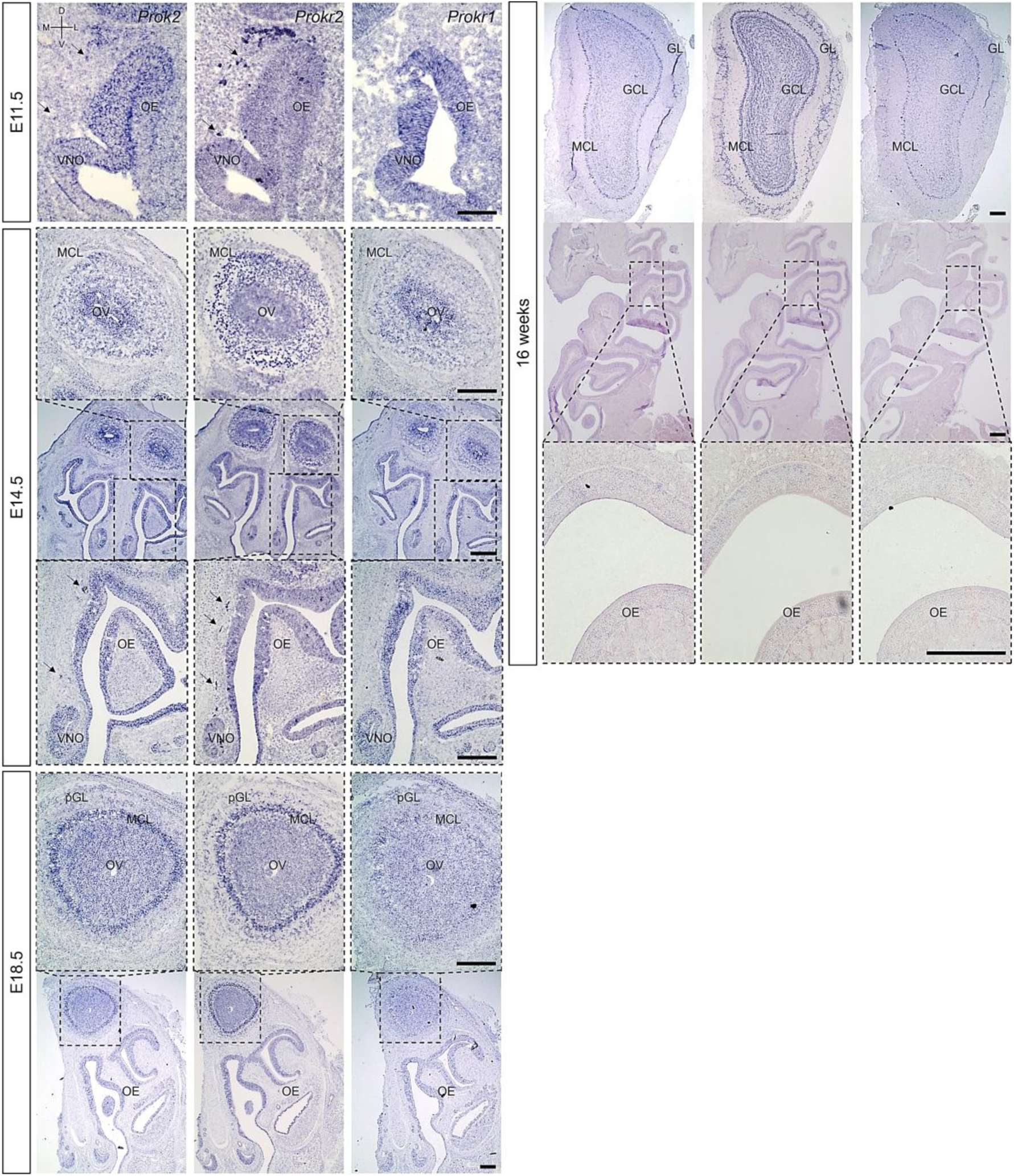
Expression of *Prok2* and its receptors in the olfactory system. Images of in situ hybridization for the olfactory system with *Prok2*, *Prokr2*, and *Prokr1* probes at E11.5. E14.5, E18.5, and 16 weeks. OE, olfactory epithelium; VNO, vomeronasal organ; OV, olfactory ventricle; MCL, mitral cell layer; (p)GL, (putative) glomerular layer; GCL, granule cell layer. D, dorsal; V, ventral; M, medial; L, lateral. Scale bar, 200 μm.

In the OB, *Prok2* was expressed in both olfactory ventricle and other cells, including the mitral cell layer (MCL) at E14.5; in the olfactory ventricle, MCL, and putative GL at E18.5; and in the MCL, a part of the GL, and a part of the granule cell layer at 16 weeks (Fig. 2). Although the overall *Prokr1* expression was similar to that of *Prok2*, it was weaker at E18.5 and at 16 weeks (Fig. 2). Notably, unlike *Prok2*, *Prokr2* was expressed at higher levels in other cells, including the MCL around the olfactory ventricle at E14.5, and was expressed overall in the GL and granule cell layer at 16 weeks (Fig. 2). This result is consistent with previous findings that *Prok2* is expressed at relatively higher levels in mature neurons, whereas *Prokr2* is expressed at relatively higher levels in immature neurons (Ng et al. 2005; Wen et al. 2019). Furthermore, the results indicate that both Prok2/Prokr2 and Prok2/Prokr1 signaling pathways play a role in the OB as well.

### Disruption of olfactory structure formation in *Prok2* KO mice during embryonic development

Olfactory structure dysgenesis in adult *Prok2* KO mice (Fig. 1A, D-G) and the expression pattern of *Prok2* and its receptor genes (Fig. 2) indicated that *Prok2* played a crucial role in olfactory structure organization during embryonic development. Accordingly, the histological changes in *Prok2* KO mice were analyzed during embryonic development. During the development of OB in WT mice, Tuj1^+^ axons projected into and penetrated the Laminin^+^ brain membrane at E11.5, the putative OB with Tuj1^+^ axons expanded, and Tuj1^+^ olfactory axons interacting and radially surrounding the OB expressed OMP, a differentiation marker of mature OSNs at E18.5 (Fig. 3A). In contrast, in KO mice, Tuj1^+^ axons stalled in front of the Laminin^+^ brain membrane from E11.5, and the Tuj1^+^ and/or OMP^+^ migratory mass, in which olfactory axons accumulated, grew depending on the developmental stage (Fig. 3A). Despite abnormal formation of the putative OB in KO embryos, OMP^+^ olfactory axons were discriminated from Nrp2^+^ vomeronasal axons among Tuj1^+^ axons in the migratory route at E18.5 (Fig. 3B). Similarly, this discrimination was also seen in the WT mice (Fig. 3B). The results indicated that GL in the OB was not constituted normally during embryonic development without *Prok2* and that homotypic fasciculation and olfactory axon maturation in the axon bundles proceeded normally regardless of OB dysgenesis. Further, OMP^+^ olfactory axons were excluded from the TH^+^ glomerular neurons and Reelin^+^ mitral cells in the putative OB of KO embryos, whereas most of these interacted with each other in the putative OB of the WT (Fig. 3C). In the abnormal GL formation in KO mice, few glomeruli, in which the OMP^+^ olfactory axons and TH^+^ glomerular neurons interacted with each other, were formed at 16 weeks but the OMP^+^ olfactory axons were excluded from putative OB rounding with TH^+^ glomerular neurons at E18.5 (Supplementary Fig. S1). This finding indicated that few glomeruli in adult *Prok2* KO mice were formed after birth.

**Figure 3.**
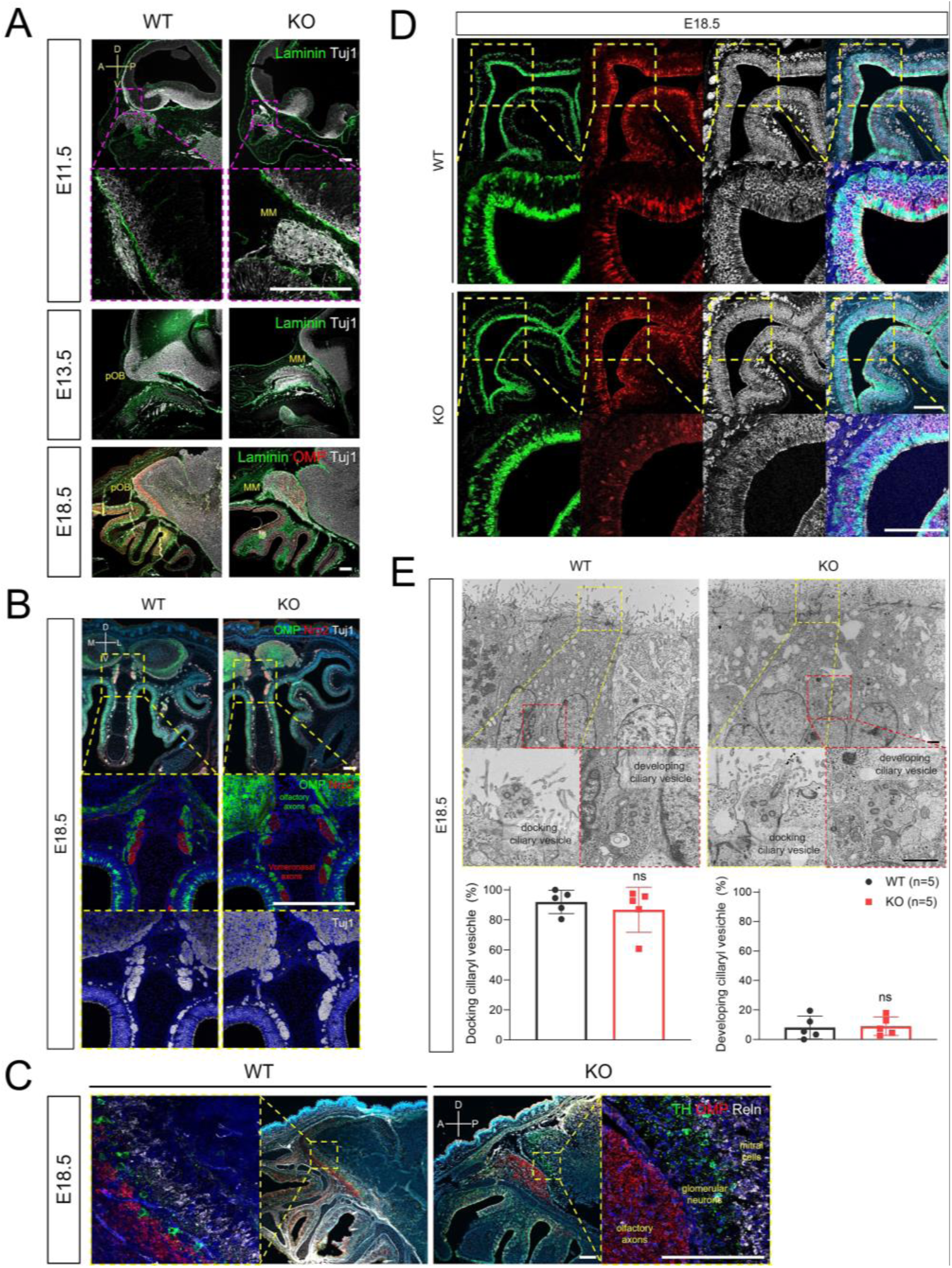
Disruption of olfactory development in *Prok2* embryo KO. Images of immunostaining and transmission electron microscopy (TEM) from WT and *Prok2* KO mice at E11.5, E14.5, and E18.5. (*A*) Immunostaining in the forebrain or pOB with α-Laminin, α-OMP, and α-Tuj1 antibodies. (*B*) Immunostaining in the olfactory nerves and vomeronasal nerves with α-OMP, α-Nrp2, and α-Tuj1 antibodies. (*C*) Immunostaining in the pOB with α-TH, α-OMP, and α-Reln antibodies. (*D*) Immunostaining in the olfactory epithelium (OE) with α-Sox2, α-OMP, and α-Tuj1 antibodies. (*E*) TEM image of olfactory vesicles and a graph of the number of olfactory vesicles. For the number of docking olfactory vesicles between WT (*n* = 5) and *Prok2* KO mice (*n* = 5) p = 0.2996 and for the number of developing olfactory vesicles between WT (*n* = 5) and *Prok2* KO mice (*n* = 5) p = 0.2904. Data presented as mean ± SD. MM, migratory mass; LP, lamina propria; pOB, putative olfactory bulb; MCL, mitral cell layer; (p)GL, (putative) glomerular layer; ONL, olfactory nerve layer. D, dorsal; V, ventral; A, anterior; P, posterior; M, medial; L, lateral. Scale bar of (*A*-*D*), 200 μm; scale bar of (*E*), 1 μm. For all panels, *n* indicates biologically independent repeats. Student’s *t*-test, p > 0.05 (ns).

In the OE, the Sox2^+^ sustentacular cells, Sox2^+^ basal cells, and Tuj1^+^OMP^−^ immature OSNs in KO mice were similar to those in the WT mice (Fig. 3D). However, the number of mature OSNs in KO mice was less than that in the WT mice (Fig. 3D). As shown in Supplementary Fig. S2, low expression of *Omp* was observed from E16.5. In the olfactory knobs of WT mice, both docking ciliary vesicles (95.18 ± 2.01%), which protrude at the OE surface, and the developing ciliary vesicles (4.82 ± 2.01%), which are located around the nuclei of sustentacular cells, were observed at E18.5 (Fig. 3E). The distribution of both docking ciliary vesicles (88.02% ± 5.65) and developing ciliary vesicles (8.41 ± 2.38%) in KO mice was not significantly different from that in WT mice (Fig. 3E). These results indicate that *Prok2* only mediates differentiation to mature OSN in the OE during embryonic development. Consequently, we found that *Prok2* played an important role in olfactory structure organization during embryonic development.

### A gene set including intermediate genes and genes expressed in the putative OB regulated by *Prok2* in olfactory development

Fewer OMP^+^ mature OSNs in the OE of *Prok2* KO mice before birth (Fig. 3D) led to a reduced proportion of mature OSNs in adults (Fig. 1E). This resulted in a reduced response against odor stimuli in the OE (Fig. 1C). To determine the underlying factor impacting decreased differentiation to mature OSNs in OE during embryonic development, the differential expression of genes in *Prok2* KO at E14.5, when *Omp* was expressed in the OE, was analyzed. The olfactory tissue of WT or KO mice was divided into the anterior and posterior regions. The anterior region included only the OE, whereas the posterior region included the OE as well as the putative OB and the other partial brain (Fig. 4A). Therefore, the former provided the information associated with only the OE, whereas the latter provided information related to the overall OE, OB, and the other partial brain. By sequencing the transcriptome of each region, the highly expressed genes (higher value of average logCPM of each gene than that in Prok2) and those expressed differentially between WT and KO mice (|fc| > 2.0, and bh.pval < 0.001) were analyzed. The results indicated that the keratin genes (*Krt77*, *Krt71*, *Krt25*) in the anterior region were significantly different between KO and WT mice (bh.pval = 7.43E-04, 7.25E-11, and 6.58E-13, respectively, in Fig. 4B and C). In the posterior region, intermediate filament genes (*Prph*, *Nefl*, *Nefm*), ion channel genes for glutamate (*Gria2*, *Grin2b*, *Slc1a2*), mitral-cell-marker genes (*Tbr1*, *Reln*), and other neuronal genes (*Nrcam*, *Nts*, *Neurod6*) were significantly different between KO and WT mice (bh.pval = 8.60E-06, 6.91.E-05, 7.31E-09, 1.02E-04, 2.38E-04, 9.31E-09, 1.75E-07, 4.32E-05, 8.22E-05, 7.25E-10 and 1.01E-06 in Fig. 4B and C, respectively). *Krt77, Prph*, *Nefl*, and *Nefm* were downregulated in KO mice, whereas the remaining genes were upregulated compared to those in the WT (Fig. 4B and C).

**Figure 4.**
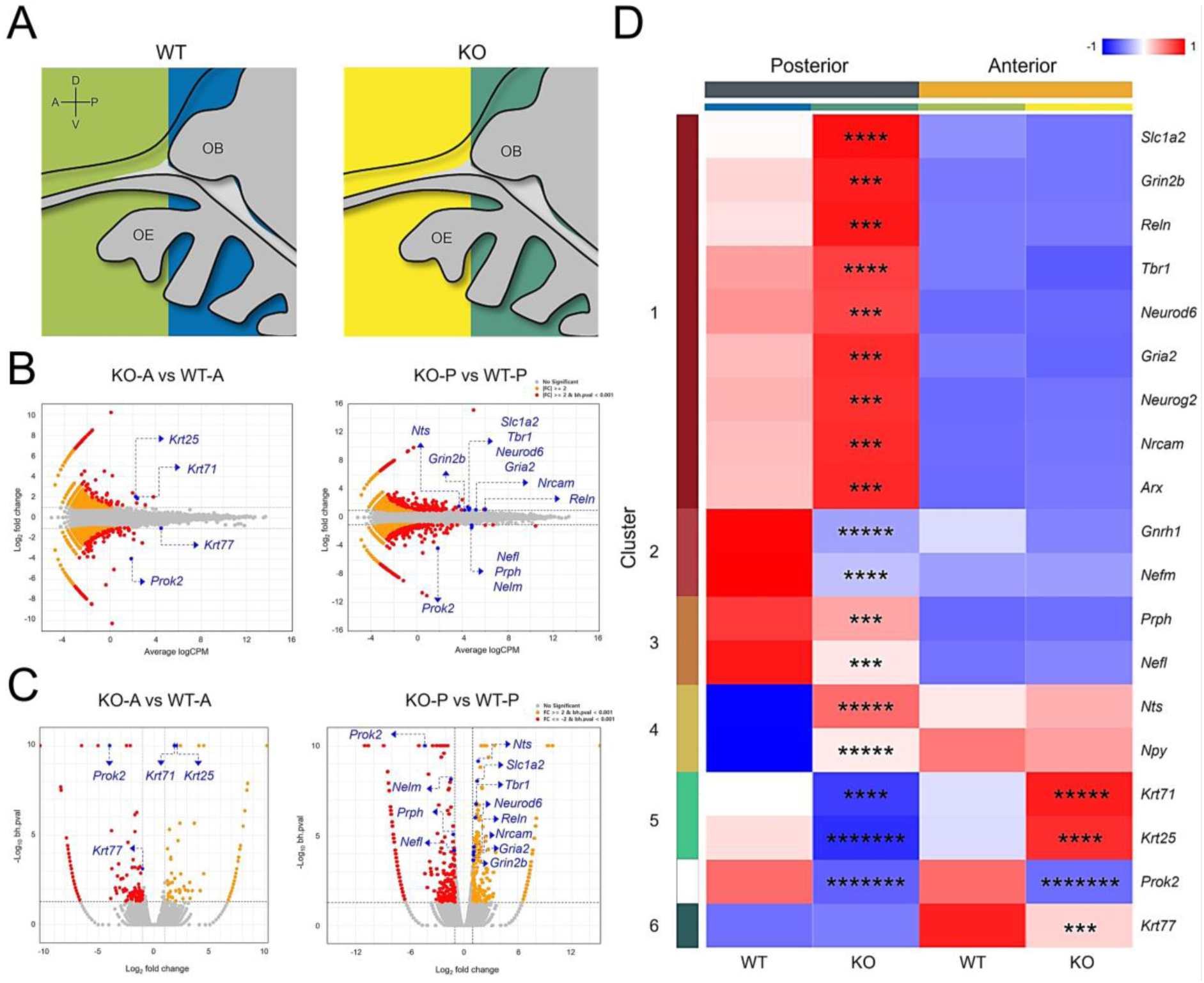
Differentially expressed genes under *Prok2* deficiency in the olfactory system at E14.5. Transcriptome sequencing of *Prok2* KO mice. (*A*) The anterior and posterior regions of the olfactory tissue. pOB, putative olfactory bulb; OE, olfactory epithelium. D, dorsal; V, ventral; A, anterior; P, posterior. (*B*) MA plot comparing KO (*n* = 3) and WT (*n* = 3) mice at each region. (C) Volcano plot comparing KO (*n* = 3) and WT (*n* = 3) mice at each region. (*D*) Heatmap comparing KO (*n* = 3) and WT (*n* = 3) mice at each region. For all panels, *n* indicates biologically independent repeats. P values were calculated using exactTest with TMM-normalized counts obtained from edgeR for statistical analysis and hierarchical clustering. The p-value was adjusted using Benjamini-Hochberg correction (bh.pval). Statistical analysis of comparison between KO and WT mice. bh.pval < 1.00E-3 (***), bh.pval < 1.00E-6 (****), bh.pval < 1.00E-9 (*****), bh.pval < 1.00E-12 (******), and bh.pval < 1.00E-15 (*******). See also Supplemental Table S1.

Similarly, the differentially expressed genes among the *Prok2*-relevant, KS-relevant, OB-constituent, OE-constituent, and olfactory receptor genes were also analyzed. Results showed that *Gnrh1* was expressed at significantly lower levels in KO mice than in the WT mice at the posterior region (bh.pval = 3.32E-11 in Supplementary Table S1). *Nrcam* and *Arx* were significantly expressed among the genes in the posterior region (bh.pval = 8.22E-05 and 3.47E-04 in Supplementary Table S1, respectively). However, none of the OE-constituent genes showed significantly different expression between KO and WT mice in any of the regions (higher value of the average logCPM for each gene than that in *Prok2* KO mice, |fold change| > 2.0, and bh.pval < 0.001 in Supplementary Table S1). Although several genes for olfactory receptors showed a higher fold change in KO mice than those in WT mice, their expression levels were substantially lower than those of other genes (average logCPM < -2.0 in Supplementary Table S1). Therefore, it is not obvious that Prok2 affected the differential expression of both OE-constituent genes and olfactory receptor genes.

A heatmap of significant differentially expressed genes in each region showed that these genes are classified into six clusters without *Prok2* (Fig. 4D). Cluster 1 included genes for glutamate receptor (*Gria2*, *Grin2b*) and glutamate transporter (*Slc1a2*), genes expressed in mitral cells or interneurons in putative OB (*Tbr1*, *Reln*, *Neurog2*, *Neurod6*, *Arx*) and an adhesion molecule gene (*Nrcam*). Clusters 2 and 3 included intermediate filament genes (*Prph*, *Nefm*, *Nefl*) along with *Gnrh1*. Cluster 4 included neuropeptide genes (*Nts* and *Npy*). Lastly, clusters 5 and 6 included keratin genes (*Krt71*, *Krt25*, *Krt77*). Clusters 1, 2, and 3 were expressed more in the posterior region than in the anterior region of the WT mice, whereas clusters 4 and 6 showed the opposite pattern. Cluster 5 was expressed similarly in both regions. These data indicate that clusters 1, 2, and 3 were expressed more in the pOB or brain than in the OE at the steady state. Clusters 4 and 6 showed the opposite pattern. Compared to those in the WT mice, clusters 1 and 4 were expressed at higher levels in the posterior region in KO mice. Conversely, clusters 2, 3, and 5 were expressed at lower levels in KO mice, whereas they were expressed in the WT mice with *Prok2* at the posterior region. The depth of the difference between the WT and KO mice in cluster 2 was greater than that in cluster 3. In the anterior region, only keratin genes were expressed significantly. Cluster 5 was expressed at higher levels in KO mice, whereas cluster 6 with *Prok2* showed the opposite pattern. Overall, the results suggested that *Prok2* mediated the development of the entire olfactory system by regulating the expression of several downstream genes at E14.5.

### Downstream genes of *Prok2* modulated OB formation during embryonic development and subsequently affected differentiation into mature OSNs in the OE

To explore how downstream genes regulated by *Prok2* organize olfactory structure during embryonic development, we observed the expression of downstream genes in olfactory structure depending on the developmental stage. Initially, we investigated where the genes noted in Fig. 4D were expressed in the olfactory system of WT mice at E14.5 (Supplementary Fig. S3). As reported previously for the expression of *Reln* and *Npy* (Ragancokova et al. 2014; Rich et al. 2018), *Reln* was expressed in the MCL, *Npy* was expressed in both MCL and olfactory nerve layer (ONL), and *Nts* was expressed in the MCL like *Reln* (thick arrows and dashed line arrows in Supplementary Fig. S3). *Slc1a2*, *Gria2*, and *Grin2b* were expressed overall in the brain with the putative OB. *Nefm* and *Nefl* were expressed in both OE and putative OB as well as in the migratory route of hormone neurons (black arrows in Supplementary Fig. S3). *Nrcam* was expressed in the NFJ and OB, similar to *Nefm* and *Nefl*, but not in the OE (black arrows in Supplementary Fig. S3). *Krt77* was expressed in the OE, putative OB, epidermis, and whisker follicles, whereas *Krt25* was expressed only in the epidermis and whisker follicles. Additionally, *Neurod6* and *Krt77* were expressed in both OE and putative OB. Based on the expression of each gene at E14.5, multi-gene expression in specific regions of the olfactory structure was explored at E11.5, E14.5, E18.5, and 16 weeks.

In OE, *Nefm*, *Nefl*, *Neurod6*, and *Krt77* were analyzed at E11.5, E14.5, E18.5, and 16 weeks together with *Omp* and *Ascl*. *OMP* expression was more dependent on the developmental stage in WT mice (Fig. 5). However, *Omp* expression in KO mice was lower than that in WT mice at E18.5 (black arrows in Fig. 5). Regardless, the other genes, including *Ascl1,* a marker of globose basal cells (GBCs), were expressed in KO mice, similar to those in the WT mice, at E14.5 and E18.5 (Fig. 5). The results suggested that *Nefm*, *Nefl*, *Neurod6*, and *Krt77* expression did not affect the suppression of *Omp* expression during embryonic development. Further, considering the arrangement of *Omp* and *Ascl1* (Fig. 5) and the cellular composition mentioned above (Fig. 1E), *Nefm* and *Nefl* were expressed in mature OSNs, immature OSNs, and basal cells, whereas *Neurod6* and *Krt77* were expressed in both sustentacular cells and GBCs of KO mice at 16 weeks (Fig. 5). These findings indicate that the cellular composition was disrupted after birth.

**Figure 5.**
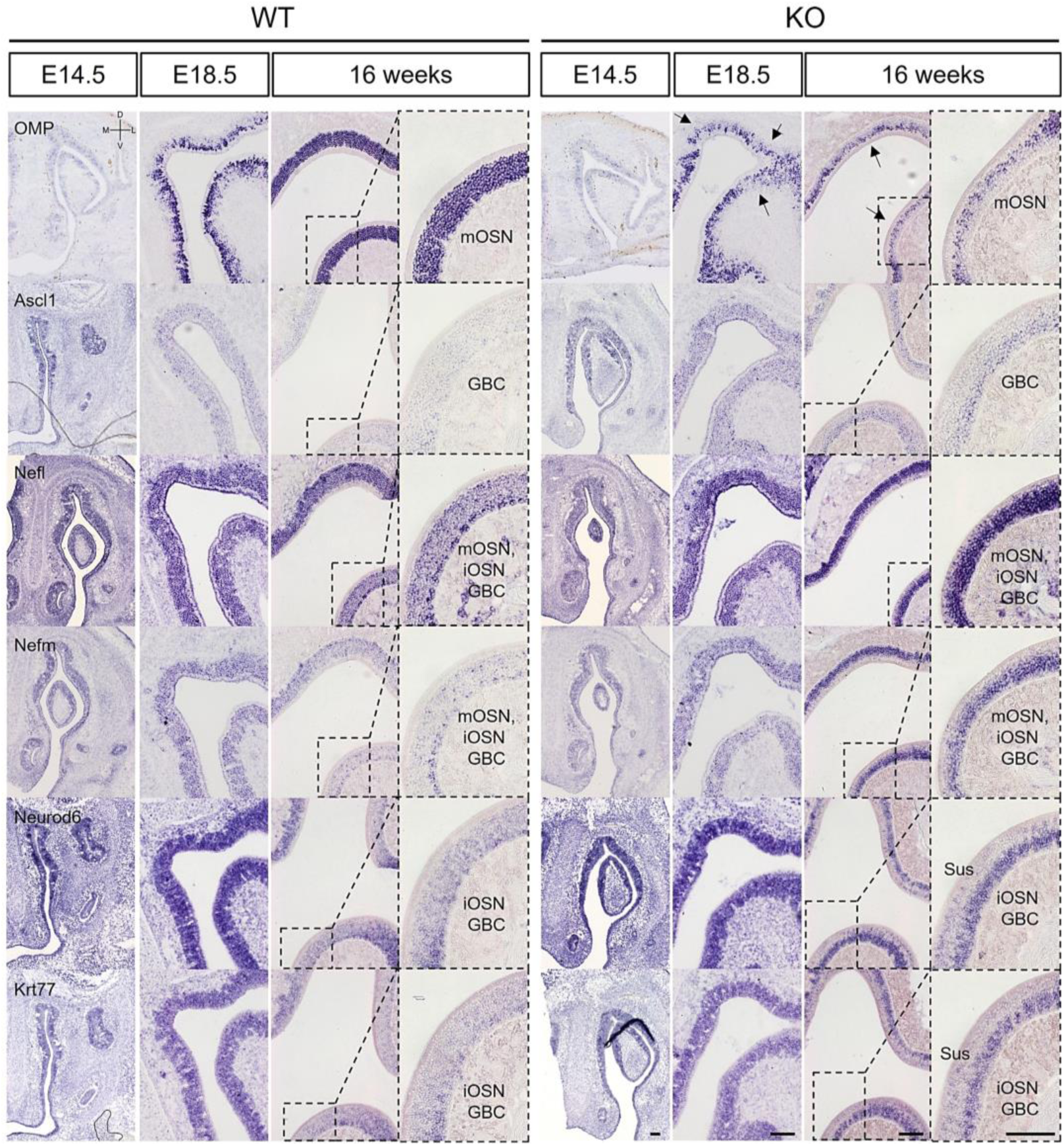
Abnormal development of the olfactory epithelium in *Prok2* KO mice at E14.5, E18.5, and 16 weeks. In situ hybridization images of the olfactory epithelium with probes for *Omp*, *Ascl1*, *Nefm*, *Nefl*, *Neurod6*, and *Krt77* at E14.5, E18.5, and 16 weeks. mOSN, mature olfactory sensory neuron; iOSN, immature olfactory sensory neuron; GBC, globose basal cell; Sus, sustentacular cells. Scale bar, 200 μm.

In the NFJ, *Nefm*, *Nefl*, and *Nrcam* expression was analyzed at E11.5 and E14.5 together with *Prokr2*. *Prokr2* was expressed near the forebrain at E11.5 in both WT and KO mice (Fig. 6). However, the spots indicating expression of *Nefm*, *Nefl*, and *Nrcam* in the NFJ of WT mice appeared cohesive in KO mice (thin arrows in Fig. 6). Furthermore, the linear expression of *Nefm* and *Nefl* in the basal region of the forebrain in WT mice was absent in KO mice (thick arrows in Fig. 6). At E14.5, the *Nefm*^+^ *Nefl*^+^ *Nrcam*^+^ spots in the NFJ of WT mice were less cohesive in KO mice (thick arrows in Fig. 6). Previous studies have reported that hormone neurons migrate from the VNO, penetrate the membrane, and cross the basal region of the brain (Schwanzel-Fukuda et al. 1994; Forni and Wray 2015; Taroc et al. 2017; Taroc et al. 2020b). Consistently, our results showed that migration was inhibited and that the migratory route at the basal region of the brain was disrupted in KO mice at E11.5. Moreover, this indicated that migration decreased in KO mice at E14.5. Overall, these results confirmed the role of *Nefm*, *Nefl*, and *Nrcam* in formation of the NFJ.

**Figure 6.**
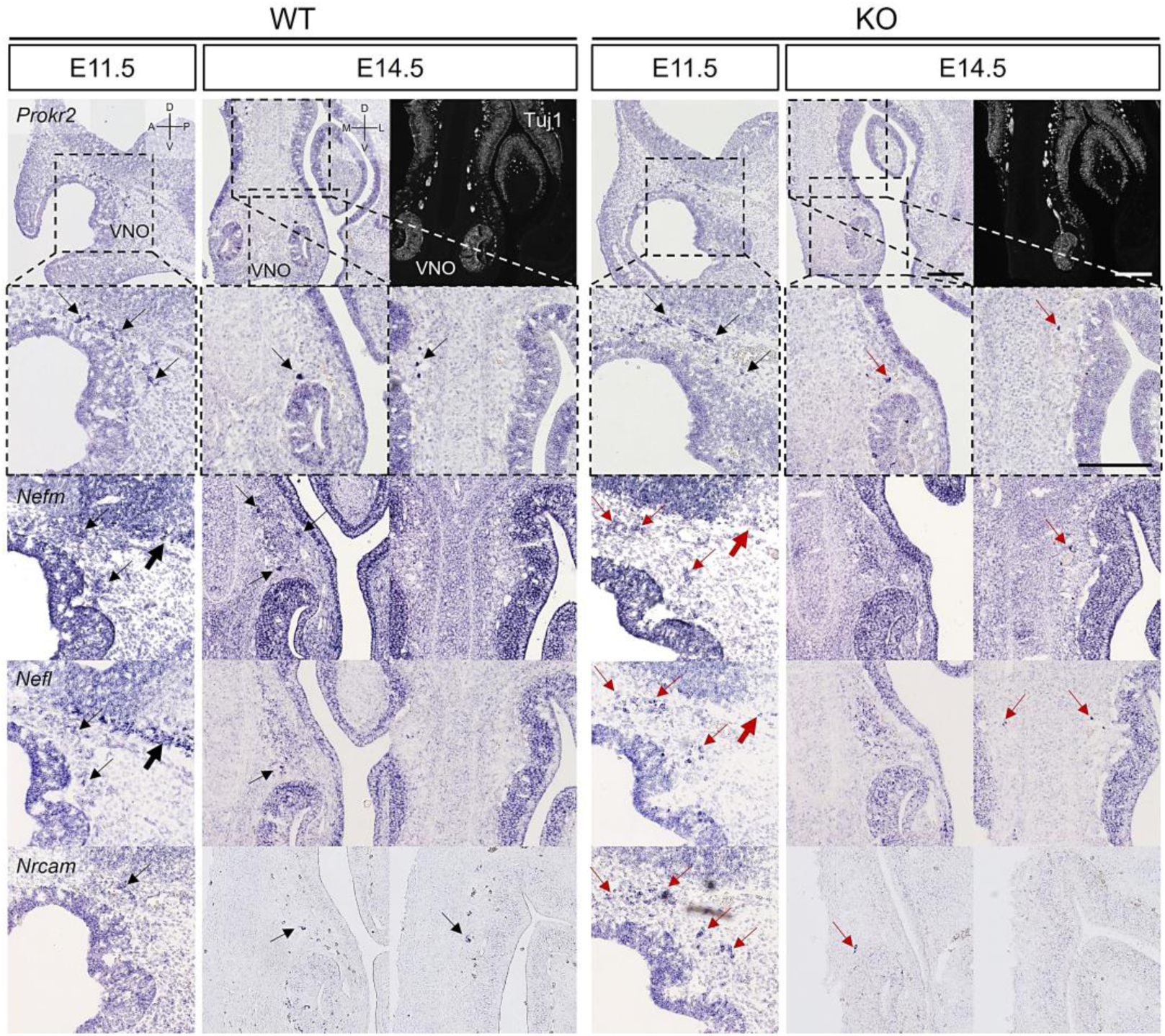
Abnormal development of the VFJ in *Prok2* KO mouse embryos at E11.5 and E14.5. In situ hybridization images of the olfactory epithelium with probes of *Prokr2*, *Nefm*, *Nefl*, and *Nrcam* at E14.5, E18.5, and 16 weeks together with IF images with α-Tuj1 antibody. VFJ, vomeronasal forebrain junction; FB, forebrain; VNO, vomeronasal organ. D, dorsal; V, ventral; A, anterior; P, posterior; M, medial; L, lateral. Scale bar, 200 μm.

The expression of *Nefm*, *Nefl*, *Nrcam*, *Neurod6*, *Slc1a2*, *Gria2*, *Grin2b*, and *Nts* was analyzed in the OB at E14.5, E18.5, and 16 weeks together with *Ascl1*. Based on *Ascl* expression, the MCL and ONL in putative OB developed normally in the WT mice (Fig. 7). In the KO mice, although the putative OB protruded similar to that in WT mice, the MCL and ONL were absent (Fig. 7). Abnormal MCL development in KO mice was also seen, as evident from the expression of *Nefm*, *Nefl*, *Slc1a2*, *Gria2*, and *Neurod6* (Fig. 7). Parallel to the absence of relevant gene expression in the basal region of the forebrain at E11.5 (thick arrows in Fig. 6), *Nefm* and *Nefl* were not notably expressed in the basal region of the putative OB (thick arrows in Fig. 7). These filament genes were expressed in the inner epidermis following the dorsal line of the rostrum in the WT mice, but not in KO mice (thin arrows in Fig. 7). Furthermore, neuronal stem cells were cohesive in KO mice but scattered in WT, as indicated by *Nrcam* expression (asterisks in Fig. 7). OB increased in size, and OB layers were specified at E18.5 and 16 weeks in WT mice (Fig. 7). As indicated by the expression levels of *Ascl1*, *Nefm*, *Nefl*, *Nrcam*, and *Slc1a2*, the protrusion of the putative OB from the subventricular zone was delayed with a bigger migratory mass and the MCL did not develop normally in the putative OB at E18.5 (Fig. 7). At 16 weeks, all layers in the OB normally developed in KO mice, but the ONL and GL interacting with OB interneurons were absent (Fig. 7). The MCL in the OB analog developed weakly and the granule cell layer was almost absent in KO mice (Fig. 7). These results suggested that *Nefm*, *Nefl*, *Nrcam*, *Neurod6*, *Slc1a2*, *Gria2*, *Grin2b*, and *Nts* affect differentiation and localization of mitral cells from neuronal stem cells and that *Nefm* and *Nefl* support the structural development of the basal brain and inner epidermis following the dorsal line of the rostrum.

**Figure 7.**
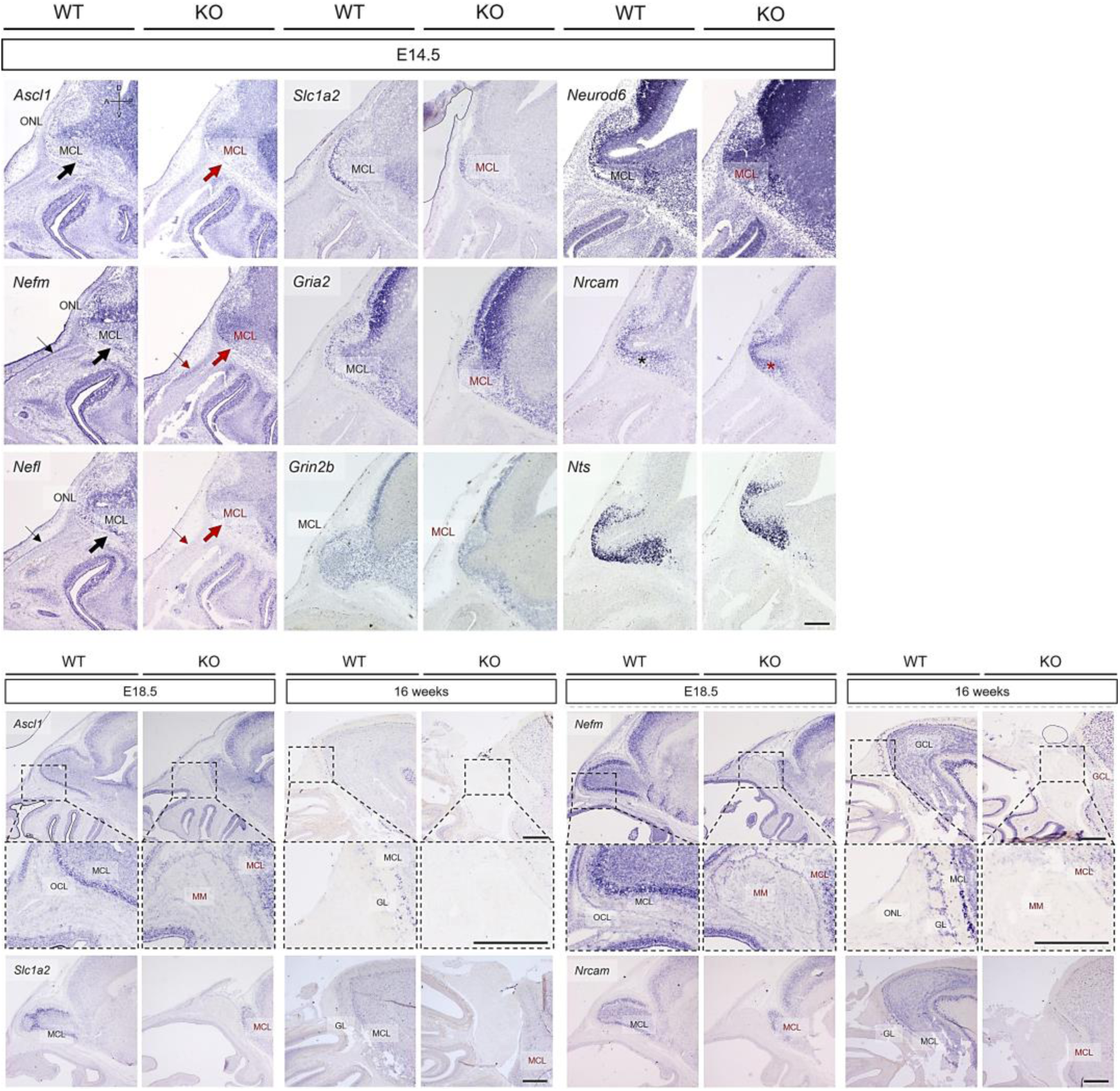
Abnormal development in the olfactory bulb of *Prok2* KO mice at E14.5, E18.5, and 16 weeks. In situ hybridization images of the olfactory bulb at E14.5, E18.5, and 16 weeks. Staining with probes for *Ascl1*, *Nefm*, *Nefl*, *Slc1a2*, *Gria2*, *Grin2b*, *Neurod6*, *Nrcam*, and *Nts*. ONL, olfactory nerve layer; MCL, mitral cell layer; MM, migratory mass; GL, glomerular layer; GCL, granule cell layer. Scale bar, 200 μm.

## Discussion

To discover how Prok2 affects the olfactory system, the processes involved in olfactory structure formation were analyzed focusing on the spatiotemporal aspects and molecular events. Overall, the intermediate filament genes and genes expressed in the putative OB induced structural changes in the olfactory system during embryonic development. Subsequently, the development of fewer mature OSNs in the OE caused olfactory dysfunction. To our knowledge, this study is the first to report the following findings: (i) Prok2/ Prokr2 signaling as well as Prok2/Prokr1 signaling are involved in olfactory development; (ii) Prok2 regulates olfactory structure and mature OSN differentiation, which is important for olfactory function; and (iii) the interaction between olfactory axons and glomerular neurons or mitral cells in the putative OB substantially contributes to mature OSN differentiation in the OE.

The main receptor of Prok2 in the olfactory system is Prokr2 (Sarfati et al. 2010; Dodé and Rondard 2013; Wen et al. 2019). However, our results showed that *Prokr1* was also expressed similar to *Prokr2* in the olfactory system except in the migration route. Thus, Prok2/Prokr1 signaling is also involved in olfactory system development. Considering the co-expression of *Prok2* and *Prokr2* in the migration route and the role of Prok2/Prokr2 signaling in forming this route, our results suggested that Prok2/Prokr1 signaling plays a role different from that of Prok2/Prokr2 signaling in the olfactory system. Future studies in a model with limited *Prokr2* expression will aid in understand the role of Prok2/Prokr1 signaling in the olfactory system.

Research on the causes of KS has primarily focused on the migration route of hormone neurons and subsequent OB formation (Schwanzel-Fukuda et al. 1994; Watanabe et al. 2009; Barraud et al. 2013; Taroc et al. 2017; Taroc et al. 2020a; Taroc et al. 2020b). These studies indicate that abnormal OB development causes congenital anosmia in KS through gene expression changes in the NFJ or OB interneurons. However, we found that olfactory dysfunction in *Prok2* KO mice resulted from defective olfactory signal detection in the OE, abnormal OE development occurred after disrupted interaction between the olfactory nerve and OB neurons during embryonic development, and intermediate filament genes and putative OB-related genes were regulated by *Prok2*. These findings highlighted that olfactory dysfunction in KS affected olfactory signaling in the OE. Furthermore, we speculated that the normal interaction between olfactory nerves and mitral cells in putative OB affected the differentiation of mature OSNs.

Several intermediate filament genes regulated by Prok2 are expressed in the olfactory system. These include keratin genes, such as Prph, which is characteristically expressed in the peripheral nervous system, and neurofilament genes. Notably, the expression of *Nefm*, *Nefl*, and *Nrcam* together with Gnrh1 was suppressed in the NFJ by the deficiency of *Prok2*. As intermediate filaments provide the cytoskeletal network by associating with adhesion molecules (Sanghvi-Shah and Weber 2017), Prok2 interacts with the migration of hormone neurons by regulating neurofilaments and neural cell adhesion molecules in the NFJ. Moreover, previous studies have reported that axonal fasciculation of homotypic nerves is manipulated by olfactory ensheathing cells and neural cell adhesion molecules in NFJ in the lamina propria before their departure to the OB (Windus et al. 2007; Ekberg et al. 2012; Barraud et al. 2013). According to these studies, the discrete separation between olfactory axons and vomeronasal axons suggested that Prok2 did not affect the role of olfactory ensheathing cells and neural cell adhesion molecules in NFJ. Consistently, our findings indicated that Prok2 did not affect the expression of *Sox10*, which regulated other genes in olfactory ensheathing cells (Supplementary Table S1). We also observed that *Krt25* and *Krt77* were expressed in the non-olfactory system similar to the epidermis and whisker follicles, indicating that Prok2 regulates the olfactory region as well as the non-olfactory region. The gene expression pattern between Krt25 and Krt77 differed because Krt77 was expressed in both olfactory and non-olfactory systems.

Moreover, neurofilament genes were even expressed in the OB. Although these genes were suppressed in the *Prok2* KO mice, cluster 1 genes shown in Fig. 4D were overexpressed in the OB. This disharmony in expression seems to be determined by whether they were expressed only in the OB. We found that penetration into the membrane of the brain and innervation with interneurons, mitral cells, and tufted cells in the OB failed in Prok2 KO mice. Moreover, owing to the defect in neuronal stem cell differentiation to projection neurons or radial glia cells at E14.5, the formation of MCL and ONL, and subsequently, the normal development of all OB layers in Prok2 KO mice failed. This abnormal development of the OB demonstrated overexpression of cluster 1 genes in the OB. Thus, our results demonstrate that Prok2 plays a role in the forebrain surrounding the olfactory nerves and in penetration of olfactory nerves into the forebrain.

Despite abnormal development in the NFJ and OB, no evident defect was seen in the OE of *Prok2* KO mice except for decreased differentiation of mature OSNs. Among olfactory receptor genes, some showed relatively higher expression levels between KO and WT mice (|fc| > 100, 1.00-E6 < bh.pval < 1.00-E2) (Supplementary Table S1). However, these olfactory receptor genes had low expression levels (average logCPM < −2) in both steady state and *Prok2* deficiency (Supplementary Table S1). Therefore, we could not accurately determine whether differentially expressed olfactory receptor genes by *Prok2* deficiency resulted in differentiation of mature OSNs. Furthermore, rather than the direct signals in the OE, indirect signals from other olfactory structures may have affected differentiation of mature OSNs.

Prior studies have reported that tufted cells such as mitral cells encountered olfactory axons and formed fascicles (Blanchart et al. 2006; Ravi et al. 2017; Tufo et al. 2022). Further, other studies showed that defasciculating of olfactory axons proceeded and that olfactory axons broke down the brain membrane in the outer ONL for sorting and refasciculating to specific glomeruli in the inner ONL (Ekberg et al. 2012; Rich et al. 2018). These findings indicate that many signals are transmitted and received between the OB and OE during the developmental process. Furthermore, our finding that abnormal organization of the MCL and ONL appeared before less differentiation of mature OSNs in the OE of *Prok2* KO mice indicated that the failure to deliver signals from the OB because of poor penetration of olfactory axons into the OB resulted in decreased differentiation of OSNs in the OE. In particular, olfactory axons did not interact with the genes expressed only in the OB, such as *Nts*, *Gria2*, and *Grin2b*. Considering that glutamate from olfactory axons is used to transfer olfactory signaling into tufted cells expressing *Gria2*, *Grin2b*, or *Slc1a2* in the OB (Gurden et al. 2006; ÒConnor and Jacob 2008), the failure of ONL and MCL formation indicates a low possibility that factors for differentiation to mature OSNs were transferred in the OE. Therefore, these genes likely affect the fundamental role in the differentiation of mature OSNs in the OE. Thus, our results indicate that Prok2 mediates the differentiation of mature OSNs via the interplay between olfactory axons and the MCL and/or ONL. Unfortunately, more direct evidence is needed to support our hypothesis regarding the role of Prok2 in differentiation of mature OSNs. Future studies using direct treatment of neurotransmitters or co-cultures with neurons or glia are required to confirm that the genes indicated in our result solely affect differentiation into mature OSNs.

Further, Prok2 may affect the cellular composition and development of olfactory cilia in the OE and GL formation in the OB, according to the interval between E18.5 and 16 weeks. These organizations are ultimately connected to each other by the generation of numerous mature OSNs. Previous studies have reported that olfactory stimulation by exposure to air after birth facilitates the generation of mature OSNs (van der Linden et al. 2020; Kim et al. 2023). Thus, the differentiation of mature OSNs during the postnatal period is dependent on olfactory stimulation. In *Prok2* KO mice, differentiation of mature OSNs during both embryonic and postnatal periods was decreased. Thus, our results suggested that Prok2 mediated the differentiation of mature OSNs through both independent and dependent olfactory stimulation. Further research on differentiation by dependent olfactory stimulation will provide valuable insights into the role of Prok2 in regulating olfactory system organization during the postnatal period. Additionally, further studies using mice models with other mutation types, such as transgenic- or knock-in mice models, can complement the limitations associated with the global deletion in *Prok2* KO mice. This approach will aid in elucidating the more specific roles of Prok2 in the organization of the olfactory system. Further, future studies may use single cell-RNA sequencing to identify the organs and cells affected by *Prok2*.

In summary, our research focused on the embryonic development of the olfactory system, specially focusing on the OE, which has been relatively unexplored. Our findings on olfactory system development provide a new insight that the OB as well as OE affect normal olfactory function and that the interaction between OE and OB is essential to each other in both development and function. Moreover, we determined the role of *Prok2* in olfactory system development based on the intermediate filament genes and genes expressed in the putative OB. Overall, these results could aid in identifying the cause of olfactory disorders and interpretation of developmental or genetic research on the olfactory system.

## Materials and Methods

### Mice

*Prok2* KO mice used in this study were purchased from UC Davis. Exons 2 and 3 in KO allele were replaced with cassettes containing LacZ and Neomycin sequences, respectively (Transcript ID: ENSMUST00000032152). Adult WT and *Prok2* KO mice aged 16 weeks (both male and female) were used in this study. The homozygous KO mice were distinguished from littermate WT mice by genotyping with the sequences: WT-F, WT-R, KO-F, and KO-R (Supplementary Table S2). Postnatal homozygous KO mice were generally acquired by natural mating. However, homozygous KO mice were acquired by in vitro fertilization between the sperms and eggs of hemizygote KO mice because they were difficult to obtain by natural mating. Embryos at E11.5, 13.5, 14.5, and 18.5 were acquired on day 11, 13, 14, and 18, respectively, after the plug was determined in virgin females. Mice were handled and maintained in strict accordance with the guidelines for the Care and Use of Laboratory Animals of Yonsei University College of Medicine. Further, the research was performed under the Yonsei Medical Center Animal Research Guidelines (IACUC No. 2023-0184), which adhere to the standards articulated in the Association for Assessment and Accreditation of Laboratory Animal Care International (AAALAC) guidelines.

### Olfactory behavior test

The olfactory behavior test used in this study is referred as an avoidance test in previous studies (Kobayakawa et al. 2007; Cho et al. 2018). Mouse tracking and measurement of time spent in each zone were automatically performed by a smart video tracking system (Smart 3.0; Harvard Apparatus©, Holliston, MA, USA). All procedures were conducted between 8 A.M. and 12 P.M., close to the dark cycle of mice. Trimethylthiazoline (TMT, purchased by BioSRQ) was used as the odorant and was diluted in water. The procedures were as follows: in the habituation step, mice were moved back and forth to the test cage for 10 min and the home cage for 5 min before proceeding to the test. The test cage had a curtain separating the cage at a 1:1 ratio, whereas the home cage was the cage in which the mouse resided. For the test, mice were placed in the zone with 20 μL of 10% TMT or water in the test cage for 3 min. The measured avoidance time in the graph was calculated by subtracting the avoidance time for TMT from the sum of avoidance time against water. For this reason, the red bar in the graph indicated the value for water.

### Electro-olfactogram (EOG)

The EOG used in this study has been previously described (Cygnar et al. 2010). Olfactory tissues were prepared from hemi-sectioned head samples and the OE was exposed in the sagittal view, immediately after sacrifice. Tissues were also maintained in the ex vivo condition in humid and warm air to detect the electrical signals analogous to those in the live animal. Amyl acetate (Sigma, #W504009) at 0.01 M and 1 M was used as the odorant. A 5–10 μm pore size pipet was hardened with 0.05% agarose and filled with Ringer’s solution (135 mM NaCl, 5 mM KCl, 1 mM CaCl_2_, 1.5 mM MgCl_2_, and 10 mM HEPES, pH 7.4, filter sterilized). The measured region in OE was the point of the quarter in the secondary turbinate. The pipet was laid on the measured region after confirming the changed voltage upon contact between the tissue and pipet. This was followed by recording of the voltage. After 0.95 s intervals, stimuli were presented in the order of vehicle, 0.01 M amyl acetate, and 1 M amyl acetate. Recording was continued for up to 10 s. The maximum amplitude in the graph was presented as the maximum of continuous values and as the mean with standard error.

### Immunostaining

Adult mice were sacrificed using CO_2_ gas, following which the olfactory tissues, including OE, OB, and partial brain, were harvested and incubated overnight in 4% PFA at 4 °C. For sampling olfactory tissues, mouse embryos were first cryoanesthetized in ice, after which their heads were harvested and incubated overnight in 4% PFA for at 4 °C. Olfactory tissues from embryos at E18.5 and adult mice were incubated for 4 days and 2 weeks, respectively, for decalcification, followed by incubation in 4% PFA for post-fixation. Slides were prepared by paraffin sectioning or cyosectioning. For paraffin sections, the procedures before the reaction for immunostaining included deparaffinization, antigen retrieval with the retrieval solution (#IW-1100; IHC World, MD, USA) in a steaming bowl (IW-1102; IHC World) for 40 min, incubation with hydrogen peroxide (#3059; Duksan, Korea) for 10 min to quench endogenous peroxidase, and washing with TBS for 5 min × 3 times sequentially. For cryosections, antigen retrieval was performed for 10 min followed by washing with TBS. After blocking with 5% BSA for 1 h, slides were incubated with primary antibodies for 1 h at RT or overnight at 4 °C. The primary antibodies used are listed in the Supplementary Table S3. After washing, the slides were washed and incubated with fluorescence-labeled secondary antibodies (Thermo Fisher Scientific, Waltham, MA, USA) for 30 min at RT. After washing, slides were mounted with medium containing DAPI (#F6057; Sigma-Aldrich, St. Louis, MO, USA).

### Transmission electron microscopy (TEM)

Olfactory tissues were fixed in Karnovsky’s fixative (2% glutaraldehyde and 2% paraformaldehyde in 0.1 M phosphate buffer, pH 7.4) for 1 day. Decalcification with 10% EDTA proceeded for 2 weeks. After washing with 0.1 M phosphate buffer, the tissues were fixed with 1% OsO4 in 0.1 M phosphate buffer for 2 h, dehydrated, and infiltrated with propylene oxide for 10 min. For preparing blocks, tissues were embedded using a Poly/Bed 812 kit (Polysciences) and polymerized in an electron microscope oven (TD-700; DOSAKA, Japan) at 65 °C for 12 h. Blocks were cut into 80 nm sections and placed on copper grids. Then, the sections were double stained with 3% uranyl acetate for 30 min and 3% lead citrate for 7 min. The stained sections were observed using a transmission electron microscope (JEM-1011; JEOL, Tokyo, Japan) at an acceleration voltage of 80 kV, equipped with a Megaview III CCD camera (Soft Imaging System, Germany). Graphs were drawn using GraphPad Prism 5.

### In situ hybridization (ISH)

For preparing probes, the sequences of each target were amplified using PCR with IP pro-Taq (#CMT2022; LaboPass) using the forward and reverse primers with the T7R sequence. The primers used in this study are presented in the Supplementary Table S2. Each tube with a mixture consisting of the PCR product, 20 U Ribonuclease inhibitors (N2111; Promega, WI, USA), 1 × DIG RNA labeling mix (#11 277 073 910; Roche, Switzerland), 1 × T7 RNA polymerase buffer, 5 mM DTT, and 50 U T7 RNA polymerase (#2540A; Takara Bio Inc., Japan) was incubated for 3 h at 37 °C. After freezing with 70% ethanol diluted with RNase-free water and 10 M LiCl for 2 h at −80 °C or overnight at −20 °C, the tube was centrifuged at 12,000 rpm for 10 min at 4 °C, the supernatant was removed, and the precipitate was dried. After elution with RNase-free water, the tube was incubated with DNase I (#04 716 728 001; Roche) for 30 min at 37 °C to remove residual DNAs. After freezing for 2 h at −80 °C or overnight at −20 °C, centrifuging for 10 min at 4 °C, and drying, the precipitation was eluted in ISH solution (5× SSC, 50% deionized formamide, 50 μg/mL heparin, 50 μg/mL yeast t-RNA, 1 mM EDTA, 0.1% CHAPS, 2% casein sodium salt, and 20% Tween 20 diluted in DEPC; pH 7.5) at a concentration of 1 μg/mL. Probes were stored at −80 °C or −20 °C.

Slides for ISH were prepared only with cryosections. The procedure for staining was as follows: before staining, slides were dried for 1 day to prevent detachment of tissues from slides. After washing with PBT (0.1% Tween 20, 1 × PBS diluted in sterile distilled water) for 5 min × 3 times, slides were bleached with H_2_O_2_ solution (#S2023; Dako) for 5 min. After washing, the slides were incubated in 10 μg/mL proteinase K solution (#PF-1048-050-02; Biosesang, Korea) for 1–5 min depending on the age of the tissue. After washing, the slides were fixed with 4% PFA for 10 min. After washing, the slides were incubated with each pre-warmed probe in ISH solution for 1 day at 68 °C. On the following day, the bound probes were stabilized by incubating the slides with pre-warmed SSC high-concentration solution (50% ionized formamide, 6 × SSC, and 1% SDS diluted in sterile distilled water) for 15 min × 3 times at 68 °C. The slides were the incubated pre-warmed low-concentration solution (50% ionized formamide and 2.4 × SSC diluted in sterile distilled water) for 15 min × 3 times at 68 °C. After washing with TBST for 5 min × 3 times at RT, slides were blocked with 5% inactivated FBS blocking solution for 1 day. Slides were then incubated overnight at 4 °C with α-DIG antibody (#11 093 274 910; Roche). After washing with TBST for 30 min × 3 times, the slides were incubated with NTMT solution for 3 × 10 min. Then, the slides were incubated for 1–2 days depending on colorization with a mixture of 250 μg/mL NBT (#11 585 029 001; Roche) and 130 μg/mL BCIP (#11 383 221 001; Roche). After incubation with 1 mM EDTA-diluted PBT for 30 min to stop the colorization reaction, slides were mounted and observed.

### RNA sequencing

After separating the heads of WT (*n* = 3) and *Prok2* KO (*n* = 3) mice at E14.5 with the nasal cavity and brain including the OB, they were divided into the anterior region or posterior region in the middle as the edge of the putative OB in the rostral view. The tissues were homogenized in TRIzol reagent (Invitrogen, #1559618), followed by RNA extraction including an on-column DNase I treatment. The cDNA library was prepared using TruSeq stranded mRNA (Illumina, USA) and sequenced using Illumina NovaSeq.

RNAseq data generated by this study have been deposited in the Gene Expression Omnibus (GEO) archive with accession number GSE277688.

### Statistics and reproducibility

GraphPad Prism (version 10.3.1) was used to test the statistical significance of the data obtained in olfactory behavior test, EOG, and TEM, and to create the graphs. The experimental results in the graphs are presented as mean with standard error (mean ± s.e.m). Statistical analyses were performed using parametric tests (two-sided unpaired t-test). Statistical significances were as follows: pval > 0.05 (ns), pval < 0.05 (*), and pval < 0.01 (**). DEG viewer system of Macrogen (Republic of Korea) was used to produce the heatmap of RNA sequence, and statistical computing and hierarchical clustering were performed using the exactTest with TMM-normalized counts obtained using edgeR. The p-value was adjusted using Benjamini-Hochberg correction (bh.pval). The levels of statistical significances were represented using asterisk(s) as follows: bh.pval < 1.00E-3 (***), bh.pval < 1.00E-6 (****), bh.pval < 1.00E-9 (*****), bh.pval < 1.00E-12 (******), and bh.pval < 1.00E-15 (*******). All phenotypic validation results were derived from at least two independent differentiations, and detailed descriptions have been provided in the corresponding methodology section. Data are presented as scatterplots, with individual data points for each measurement. All experiments were conducted using at least three biological replicates from different independent experiments. The exact number of biological replicates and independent experiments is reported in the respective figure legends.

## Acknowledgments

This research was supported by the Bio & Medical Technology Development Program of the National Research Foundation (NRF) & funded by the Korean government (MSIT) (No. RS-2024-00443043) and by the National Research Foundation of Korea (NRF) grant funded by the Korea government (MSIT) (No. RS-2024-00344509).

## Author Contributions

B.K.: contributed to conceptualization, methodology, analysis, and writing; M.R., H.J., J.Y., and C.K.: contributed to investigation, supervision, and review editing. All authors have read and agreed to the published version of the manuscript.

